# Differential activity and expression of human 5β-reductase (AKR1D1) splice variants

**DOI:** 10.1101/2020.06.09.142539

**Authors:** Nathan Appanna, Elena Gangitano, Niall J Dempster, Karen Morris, Sherly George, Brian G Keevil, Trevor M Penning, Laura L Gathercole, Jeremy W Tomlinson, Nikolaos Nikolaou

**Affiliations:** Oxford Centre for Diabetes, Endocrinology and Metabolism, NIHR Oxford Biomedical Research Centre, University of Oxford, Churchill Hospital, Oxford, UK, OX3 7LE, UK; Department of Experimental Medicine, Sapienza University of Rome, Rome, Italy; Biochemistry Department, Manchester University NHS Trust, Manchester Academic Health Science Centre, Manchester, M23 9LT, UK; Center of Excellence in Environmental Toxicology and Department of Systems Pharmacology & Translational Therapeutics, University of Pennsylvania Perelman School of Medicine, 1315 BRB II/III 421 Curie Blvd, Philadelphia, PA 19104-6160; Department of Biological and Medical Sciences, Oxford Brookes University, Oxford, OX3 0BP, UK

**Keywords:** steroids, dexamethasone, cortisol, liver, testis, metabolism

## Abstract

Steroid hormones, including glucocorticoids and androgens, exert a wide variety of effects in the body across almost all tissues. The steroid A-ring 5β-reductase (AKR1D1) is expressed in human liver and testes, and three splice variants have been identified (*AKR1D1-001, AKR1D1-002, AKR1D1-006*). Amongst these, *AKR1D1-002* is the best described; it modulates steroid hormone availability and catalyses an important step in bile acid synthesis. However, specific activity and expression of *AKR1D1-001* and *AKR1D1-006* are unknown.

*AKR1D1-002, AKR1D1-001* and *AKR1D1-006* were measured in human liver biopsies and human hepatoma cell lines by qPCR. Three-dimensional (3D) structures of *AKR1D1* variants were determined using *in silico* approaches. *AKR1D1* variants were over-expressed in HEK293 cells, and successful overexpression confirmed by qPCR and western blotting. Steroid hormone clearance was measured by mass spectrometry and ELISA, and steroid receptor activation determined by luciferase reporter assays.

*AKR1D1-002* and *AKR1D1-001* are expressed in human liver, and only *AKR1D1-006* is expressed in human testes. Following over-expression in HEK293 cells, AKR1D1-001 and AKR1D1-006 protein levels were lower than AKR1D1-002, but significantly increased following treatment with the proteasomal inhibitor, MG-132. AKR1D1-002 efficiently metabolised glucocorticoids and androgens and decreased receptor activation. AKR1D1-001 and AKR1D1-006 poorly metabolised dexamethasone, but neither protein metabolised cortisol, prednisolone or testosterone.

We have demonstrated the differential expression and role of *AKR1D1* splice variants to regulate steroid hormone clearance and receptor activation. AKR1D1-002 is the predominant functional protein in steroidogenic and metabolic tissues. In addition, AKR1D1-001 and AKR1D1-006 may have a limited role in the regulation of synthetic glucocorticoid action.

## 1. Introduction

Steroid hormones, including glucocorticoids, androgens and oestrogens, are fat-soluble molecules synthesised from cholesterol that play a crucial role in development, differentiation and metabolism (Simons 2008). Glucocorticoids, produced by the adrenal cortex, are released in response to stress and, following binding to their cognate receptor, the glucocorticoid receptor (GR), regulate anti-inflammatory and metabolic processes. Androgens are predominantly produced by the male testes, but also by the adrenal glands and the ovaries in females. Upon binding to the androgen receptor (AR), they have multiple actions, including the initiation of adrenarche and stimulation and control of secondary sexual characteristics. Following synthesis and delivery into the circulation, steroid hormones can be reduced, oxidised or hydroxylated by a variety of enzymes, including the 11β-hydroxysteroid dehydrogenases (11β-HSD) and the A-ring reductases (5α-reductases, [5αR] and 5β-reductase [AKR1D1]) in a tissue-specific manner. Dysregulation of steroid hormone levels and their metabolism has been associated with adverse metabolic features, including insulin resistance, hypertension, glucose intolerance and hepatic triacylglycerol (TG) accumulation (Morton *et al.* 2001; Wang 2005; Tomlinson *et al.* 2008; Nasiri *et al.* 2015; Navarro *et al.* 2015). The steroid 5β-reductase is encoded by the *AKR1D1* (aldo-keto reductase 1D1) gene, and is highly expressed in the liver, where it inactivates steroid hormones, including glucocorticoids and androgens, and catalyses a fundamental step in bile acid synthesis (Chen *et al.* 2011; Jin *et al.* 2014). AKR1D1 utilises NADPH as the hydride donor and catalyses a stereospecific irreversible double bond reduction between the C4 and C5 positions of the A-ring of steroids. 5β-reduction of steroids is unique in steroid enzymology as it introduces a 90° bend and creates an A/B *cis*-ring junction, resulting in the formation of steroids with different properties from either the α,β-unsaturated or 5α-reduced steroids (which possess a largely planar steroidal structure with an A/B *trans* ring-junction) (Jin *et al.* 2014). The human AKR1D1 protein is highly homologous with other members of the AKR1 family, including the AKR1C subfamily (which includes the hydroxysteroid dehydrogenases), the AKR1A subfamily (aldehyde reductases) and the AKR1B subfamily (aldose reductases) (Jez *et al.* 1997). The human gene for *AKR1D1* consists of 9 exons, and three splice variants have been identified, all of which are predicted to lead to functional proteins: AKR1D1-001 (*NM_001190906*), AKR1D1-006 (*NM_001190907*), and AKR1D1-002 (*NM_005989*) (Barski *et al.* 2013). AKR1D1-002 encodes a 326 amino acid 5β-reductase enzyme that includes all 9 exons. AKR1D1-001 lacks exon 5 and is translated into a 285 amino acid protein, whilst, AKR1D1-006 omits exon 8 and translates into a 290 amino acid protein (Barski *et al.* 2013) (Fig. 1). Loss of function mutations in the *AKR1D1* gene have been reported in patients with 5β-reductase deficiency, and are associated with decreased 5β-reduced urinary corticosteroids and impaired bile acid synthesis (Palermo *et al.* 2008).

**Figure 1.**
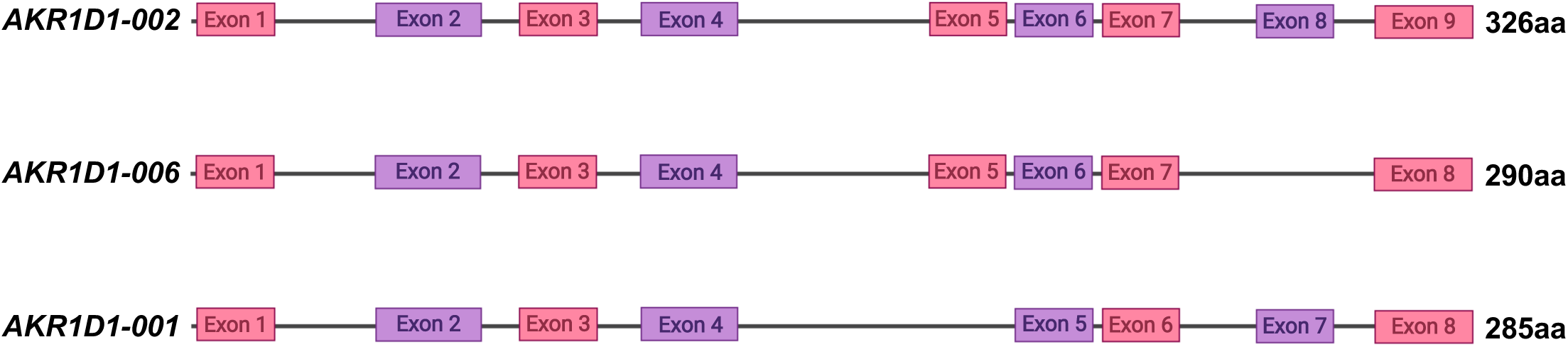
(A) AKR1D1 splice variants, showing the full form of *AKR1D1-002*, resulting in a 326 amino acid (aa) protein, as well as *AKR1D1-001* and *AKR1D1-006* that lack exons 5 and 8, resulting in 285 and 290 amino acid proteins, respectively.

Amongst all *AKR1D1* splice variants (*AKR1D1*-SVs), *AKR1D1-002* is the best characterised and represents the full-length wild type protein. In both cellular and cell-free system models, AKR1D1-002 metabolises endogenous glucocorticoids, including cortisol and cortisone, to their 5β-reduced metabolites, 5β-dihydrocortisol and 5β-dihydrocortisone, respectively. The 5β-reduced metabolites are then converted, in a non-rate limiting step, to their inactive tetrahydro-metabolites (5β-tetrahydrocortisol and 5β-tetrahydrocortisone) by the hepatic 3α-hydroxysteroid dehydrogenases, AKR1C1-C4, with downstream effects on steroid receptor activation and target gene transcription (Chen *et al.* 2011; Nikolaou *et al.* 2019a). Crucially, AKR1D1-002 is also implicated in drug metabolism, as it metabolises the synthetic glucocorticoids prednisolone and dexamethasone, leading to the formation of inactive 5β-reduced products (Nikolaou *et al.* 2020).

The relative expression levels of *AKR1D1*-SVs in human tissues have not been identified. Similarly, it is entirely unexplored whether the truncated *AKR1D1*-SVs lead to functional proteins, and whether they have a role in glucocorticoid, androgen and drug metabolism. The aim of our study was therefore to define the ability of *AKR1D1*-SVs to regulate endogenous and synthetic steroid availability in appropriate human cellular models, as well as to determine the expression levels of *AKR1D1*-SVs in human cells.

## 2. Materials and methods

### 2.1. Genotype-Tissue expression data (GTEx)

*AKR1D1* splice variant expression data (ENSG00000122787.14) were extracted from the Genotype-Tissue Expression (GTEx) Project (https://www.gtexportal.org). GTEx was supported by the Common Fund (https://commonfund.nih.gov/GTEx) of the Office of the Director of the National Institutes of Health, and by NCI, NHGRI, NHLBI, NIDA, NIMH, and NINDS. The data used for the analyses described in this manuscript were obtained from dbGaP Accession phs000424.v8.p2 on 24/04/2020.

### 2.2. Cell culture and human liver tissue

Liver biopsy samples originated from the Oxford Gastrointestinal Illness Biobank (REC reference 16/YH/0247). HepG2 cells (Cat#HB-8065) and HEK293 cells (Cat#CRL-11268) were purchased from ATCC. Huh7 cells were purchased from the Japanese Cancer Research Resources Bank (Cat#JCRB0403). All cell lines were cultured in Dulbecco’s Minimum Essential Medium (DMEM) (Thermo Fisher Scientific, Massachusetts, USA), containing 4.5 g/L glucose, and supplemented with 10% foetal bovine serum, 1% penicillin/streptomycin and 1% non-essential amino acids (Thermo Fisher Scientific, Massachusetts, USA). Cells were grown at 37°C in a 5% CO_2_ setting.

Dexamethasone, cortisol, prednisolone, testosterone and MG-132 were purchased from Sigma-Aldrich (Dorset, UK). For all cell treatments, HEK293 cells were cultured in serum-free and phenol red-free media containing 4.5 g/L glucose (Sigma-Aldrich, Dorset, UK), 1% penicillin/streptomycin, 1% non-essential amino acids and 1% l-glutamine (Sigma-Aldrich, Dorset, UK).

### 2.3. Transfection studies

For over-expression transfection studies, 1×10^5^ cells/well were plated in 24-well Cell Bind plates (CORNING) 24 hours prior to transfection with either empty pCMV6 vector (#PCMV6XL4), pCMV6+AKR1D1-002 (#SC116410), pCMV6+AKR1D1-001 (#RC231126) or pCMV6+AKR1D1-006 (#RC231133) construct variants (Origene Technologies, Rockville, USA). 0.5 µg DNA construct and 1 µL X-tremeGENE DNA Transfection Reagent (Roche, Hertfordshire, UK) were diluted in 50 µL OPTIMEM serum-free media (Invitrogen). The mixture was vortexed and incubated at room temperature for 20 min; 50 µL was added to each well and cells incubated at 37°C for 48 hours before treatment.

For cell treatments, HEK293 cells were cultured in serum-free and phenol red-free media containing 4.5 g/L glucose, and either steroid hormone for 24 hours, post-transfection. For proteasome inhibition studies, cells were cultured in serum-free media, and 20 µM MG-132 were added 3 and 6 hours prior to harvesting.

### 2.4. RNA extraction and gene expression

Total RNA was extracted from cells using the Tri-Reagent system (Sigma-Aldrich, Dorset, UK) and RNA concentrations were determined spectrophotometrically at OD260 on a Nanodrop spectrophotometer (ThermoFisher Scientific, Massachusetts, USA). Reverse transcription was performed in a 20 μL volume; 1 μg of total RNA was incubated with 10x RT Buffer, 100 mM dNTP Mix, 10x RT Random Primers, 50 U/μL MultiScribe Reverse Transcriptase and 20 U/μL RNase Inhibitor (ThermoFisher Scientific, Massachusetts, USA). The reaction was performed under the following conditions; 25°C for 10 min, 37°C for 120 min and then terminated by heating to 85°C for 5 min.

All quantitative PCR (qPCR) experiments were conducted using an ABI 7900HT sequence detection system (Perkin-Elmer Applied Biosystems, Warrington, UK). Reactions were performed in 6 μL volumes on 384-well plates in reaction buffer containing 3 μL of 2 x Kapa Probe Fast qPCR Master Mix (Sigma-Aldrich, Dorset, UK). All probes were supplied by Thermo Fisher Scientific (Massachusetts, USA) as predesigned FAM dye-labelled TaqMan Gene Expression Assays (*AKR1D1* spanning exons 3-4: Hs00973526_g1, *AKR1D1* spanning exons 5-6: Hs00973528_gH, *AKR1D1* spanning exons 7-8: Hs00975611_m1). The reaction conditions were; 95°C for 3 min, then 40 cycles of 95°C for 3 sec and 60°C for 20 sec. The Ct (dCt) of each sample using the following calculation (where E is reaction efficiency – determined from a standard curve): ΔCt = E[min Ct-sample Ct] using the 1/40 dilution from a standard curve generated from a pool of all cDNAs as the calibrator for all samples. The relative expression ratio was calculated using the following: Ratio= ΔCt[target]/ ΔCt[ref] and expression values were normalized to *18SrRNA* (Hs03003631_g1), *TBP* (Hs00427620_m1) and *ACTB* (Hs01060665_g1) (Pfaffl 2001).

### 2.5. Luciferase reporter assay

To determine AR activation, HEK293 cells were plated in 24-well Cell Bind plates (CORNING, Flintshire, UK) and co-transfected with either empty pCMV6 vector (#PCMV6XL4), pCMV6+AKR1D1-002, pCMV6+AKR1D1-001 or pCMV6+006 construct variants, followed by treatments with serum-free and phenol-red free cell media containing testosterone for a further 24 hours. Cell media aliquots (500 µL) were collected and stored at −20oC. In another set of experiments, HEK293 cells were transiently co-transfected with a pcDNA3.1+AR construct and an androgen responsive element (ARE) reporter - a mixture of an inducible ARE-responsive firefly luciferase construct and a constitutively expressing renilla luciferase construct (#CCS-1019L, QIAgen, Manchester, UK). 48 hours post-transfection, cell media was replaced with the steroid containing media aliquots described above, and cells were incubated for 24 hours. Cell lysates were harvested in passive lysis buffer, and reporter activity was measured using the Luciferase Assay System (Promega, Wisconsin, USA) and an EnSpire Multimode plater reader (PerkinElmer, Massachusetts, USA). The data were presented as the % ratio of firefly to renilla luciferase activity (Fluc/Rluc).

### 2.6. Protein extraction and immunoblotting

Cells were lysed using RIPA buffer (Sigma-Aldrich, Dorset, UK), supplemented with protease and phosphatase inhibitor cocktails (both 1/100) (Thermo Fisher Scientific, Massachusetts, USA). Protein concentrations were determined using the Bio-Rad protein assay (Bio-Rad Laboratories Inc., Hercules, CA), according to the manufacturer’s instructions. Primary human AKR1D1 (dilution 1/250 - HPA057002, Atlas Antibodies AB, Bromma, Sweden), β-tubulin (#15115, monoclonal) (Cell Signalling, Danvers, USA), β-actin (#3700, monoclonal) (Cell Signalling, Danvers, USA), and secondary antibodies (P044801-2, polyclonal) from Dako (Agilent, Santa Clara, USA) were used at a dilution 1/1000 (primary) and 1/2000 (secondary) respectively, unless stated otherwise. Bands were visualised with Bio Rad Clarity Western ECL (Watford, Hertfordshire, UK) and ChemiDocXS imager (Bio Rad, Watford, Hertfordshire, UK). Western blots were quantified by densitometry analysis using ImageJ (NIH, Bethesda, MD, http://rsb.info.nih.gov/ij), normalised to β-tubulin and β-actin to correct for variability in gel loading.

### 2.7. Steroid hormone measurements

Cell media, cortisol, prednisolone and dexamethasone concentrations were measured using quantitative liquid chromatography–mass spectrometry (LC-MS/MS) in selected ion-monitoring analysis using previously published methods (Owen *et al.* 2005; Hawley *et al.* 2018). The lower limit of quantitation was 5.2 nmol/L for prednisolone, 0.25 nmol/L for dexamethasone, and 22 nmol/L for cortisol.

Cell media testosterone levels were determined using a commercially available testosterone ELISA assay according to the manufacturer’s protocol (#ab108666, Abcam, Cambridge, UK).

### 2.8. Statistics

Data are presented as mean ± standard error (se), unless otherwise stated. Normal distribution was confirmed using Shapiro-Wilk test. Two-tailed, paired t-tests were used to compare single treatments to control. For comparisons between control and different treatments, statistical analysis was performed using one-way analysis of variance (ANOVA) with Dunnett corrections. Statistical analysis on qPCR data was performed on mean relative expression ratio values (Ratio= ΔCt[target]/ ΔCt (Pfaffl 2001)). Data analysis was performed using Graphpad Prism software (Graphpad Software Inc, La Jolla, USA) and considered statistically significant at p<0.05, unless otherwise stated.

## 3. Results

### 3.1. *AKR1D1*-SVs *are differentially expressed in human liver and testes*

To define the transcription levels of *AKR1D1* in human tissues, data were initially extracted from publicly available databases (https://www.gtexportal.org). Consistent with previous studies, AKR1D1 is predominantly expressed in the liver [median transcripts per million (TPM)=51.66; n=226], although significant levels of expression have also been detected in human testes (median TPM=4.87; n=381) (Fig. 2A). Low levels of *AKR1D1* have also been detected in human adrenal glands (median TPM=0.07; n=258) and mammary breast (median TPM=0.02; n=459). No transcription levels were however detected in other metabolic or steroidogenic tissues, including adipose [subcutaneous (n=663) or visceral (n=541)], skeletal muscle or ovary (n=180) (Fig. 2A). Specifically with regards to *AKR1D1*-SVs, database analysis revealed *AKR1D1-002* as the predominant transcript expressed in the liver, with lower levels of *AKR1D1-001* expression, and almost no detectable *AKR1D1-006* expression (Fig. 2B).

**Figure 2.**
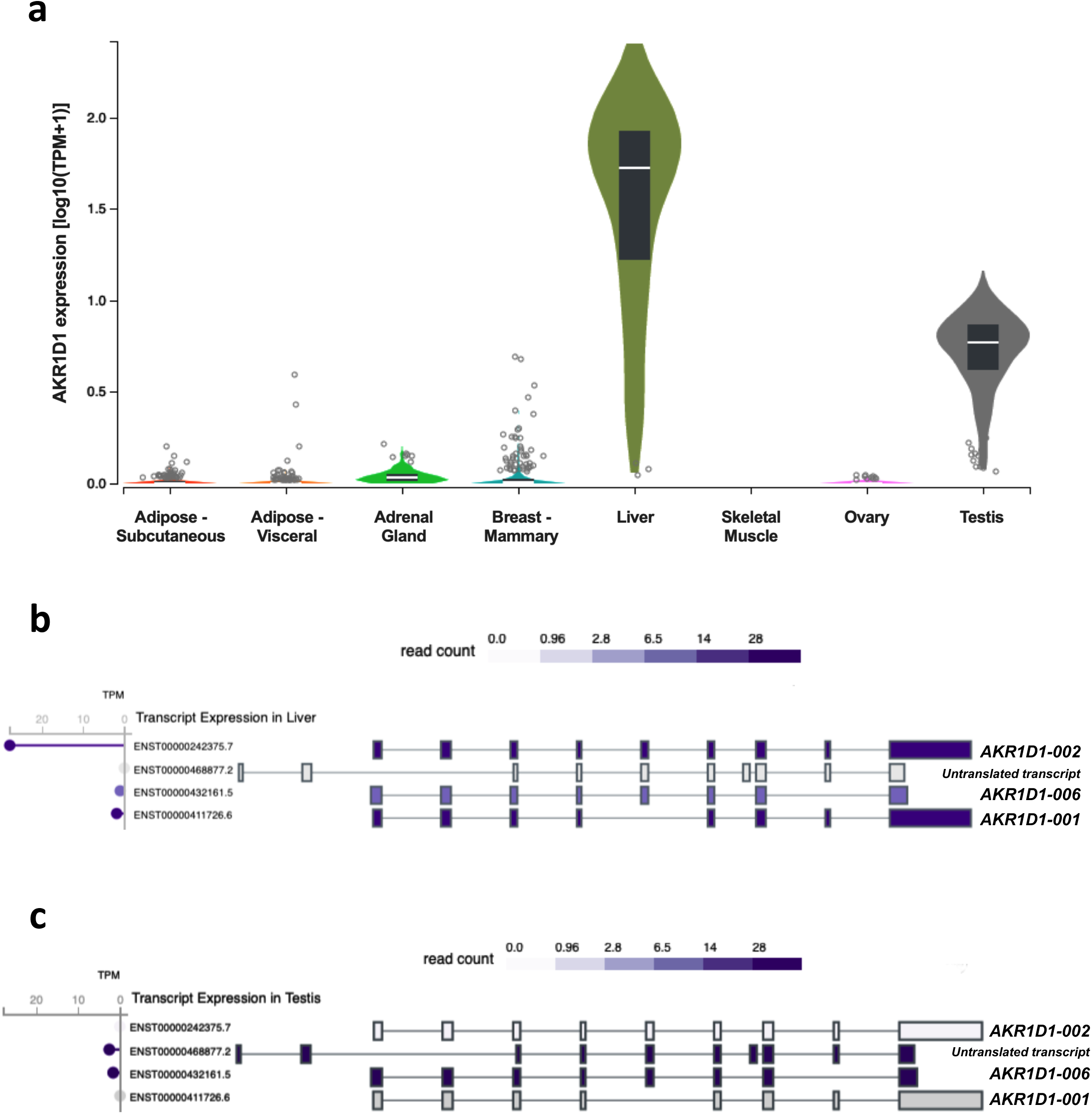
(A) *AKR1D1* transcript levels in human adipose, adrenal gland, breast, liver, skeletal muscle, ovary and testicular tissues^$^. (B) Transcript expression levels of AKR1D1 splice variants in human liver^$^. (C) Transcript expression levels of AKR1D1 splice variants in human testes^$^. ^$^Data extracted from https://www.gtexportal.org (GTEx). GTEx-extracted expression values are shown as transcripts per million (TPM). Box plots are shown as median and 25^th^ and 75^th^ percentiles. Points displayed are outliers 1.5 times above or below the interquartile range.

Additional database analysis exploring testicular *AKR1D1* expression demonstrated that the *AKR1D1-006* splice variant was the main transcript present, with no expression of either *AKR1D1-002* or *AKR1D1-001* (Fig. 2C). Interestingly, a fourth transcript variant (ENST00000468877.2) was also detected in human testes, however, this variant is not translated due to the lack of an opening reading frame within its sequence.

### 3.2. *AKR1D1*-SVs *are variably expressed in human liver and are differentially targeted for proteasomal degradation*

*AKR1D1* variant-specific expression was confirmed using multiple pre-designed TaqMan gene expression assay systems. Firstly, an assay spanning the 3-4 exon-exon junction in the *AKR1D1* mRNA was used to identify all variants. A second TaqMan assay spanning the 5-6 exon-exon junction was able to identify *AKR1D1-002* and *-006*, but not -*001* transcripts. Finally, a third assay was used spanning the 7-8 exon-exon junction that was able to identify *AKR1D1-002* and *-001*, but not *-006* transcripts (Fig. 3A).

**Figure 3:**
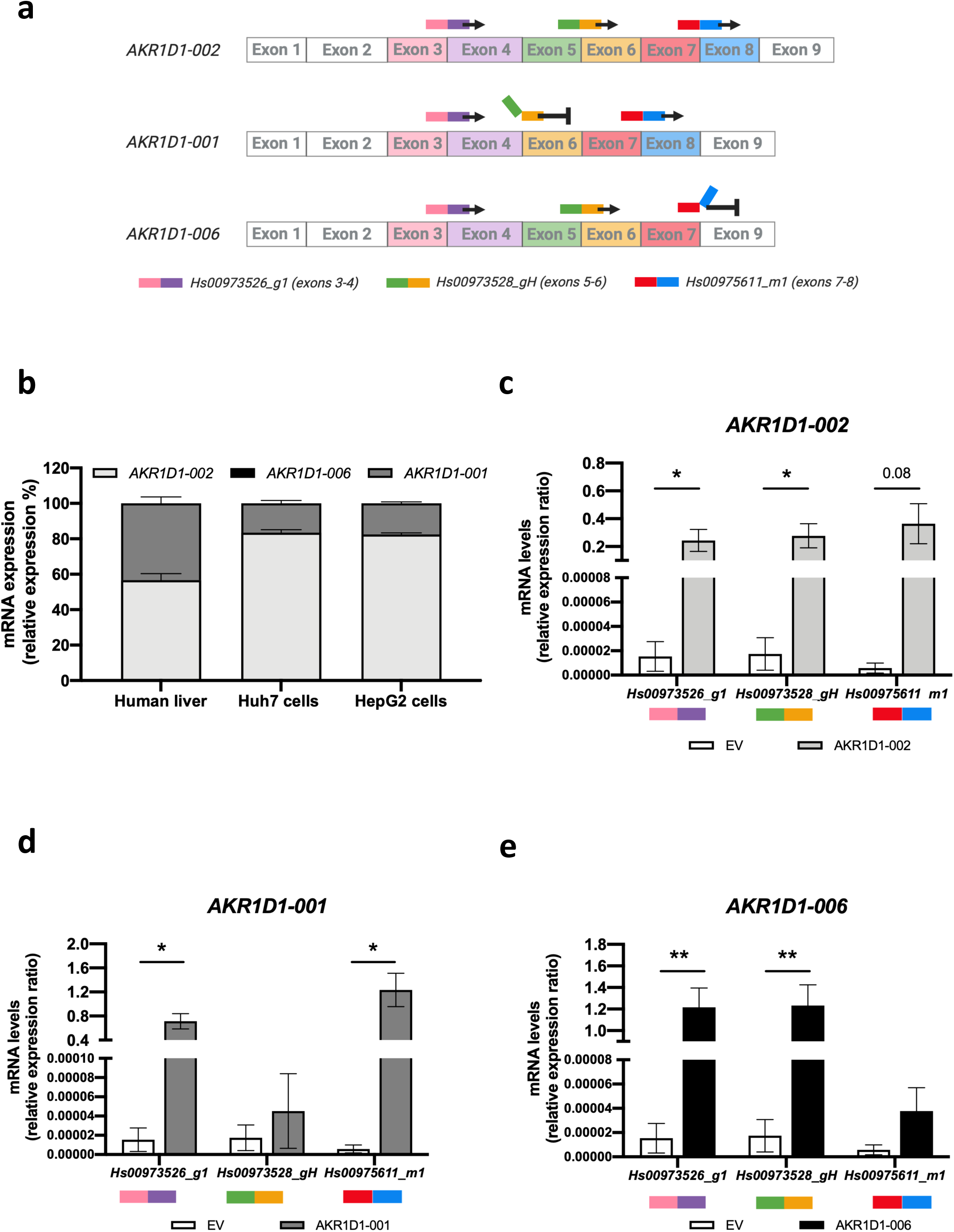
(A) Schematic representation demonstrating the mechanism of TaqMan qPCR targeting the different *AKR1D1* splice variants using specific exon-exon junction primers. (B) Relative mRNA expression levels of *AKR1D1-001, AKR1D1-002* and *AKR1D1-006* splice variants in male and female human liver biopsies (n=8), Huh7 (n=4) and HepG2 (n=4) hepatoma cell lines. mRNA over-expression levels of (C) *AKR1D1-002*, (D) *AKR1D1-001*, and (E) *AKR1D1-006* in HEK293 cells, confirmed using multiple exon-exon junction Taqman primer assays (n=4). Human liver qPCR data were normalised to *18SrRNA, TBP* and *ACTB*, and cell line data were normalised to *18SrRNA*. mRNA expression data are presented as mean±se, performed in triplicate.

qPCR analysis in liver biopsies from male and female donors (n=8) revealed that *AKR1D1-002* is the most highly expressed splice variant in human liver (56.7±3.6%), followed by *AKR1D1-001* (43.3±3.6%), and a complete lack of *AKR1D1-006* expression. Similarly, in HepG2 and Huh7 cells, *AKR1D1-002* was the predominant splice variant (HepG2: 82.5±0.9%; Huh7: 83.5±1.6%), with lower levels of *AKR1D1-001* (HepG2: 17.5±0.8%; Huh7:16.5±1.6%), and no *AKR1D1-006* expression (Fig.3B).

To define the functional role of *AKR1D1*-SVs in the regulation of steroid hormone metabolism *in vitro*, HEK293 cells were transfected with either empty pCMV6 (EV), *AKR1D1-002, AKR1D1-001, or AKR1D1-006* containing vectors for 48 hours. Successful over-expression of each SV was confirmed using the expression system assays described above (Fig 3C, D and E).

*AKR1D1* over-expression was confirmed at protein level by western blotting. Although protein expression levels of all *AKR1D1*-SVs were detected, there were significantly lower levels of AKR1D1-001 and AKR1D1-006 expression, compared to AKR1D1-002, indicative of rapid intracellular proteasomal degradation (Fig. 4A).

**Figure 4:**
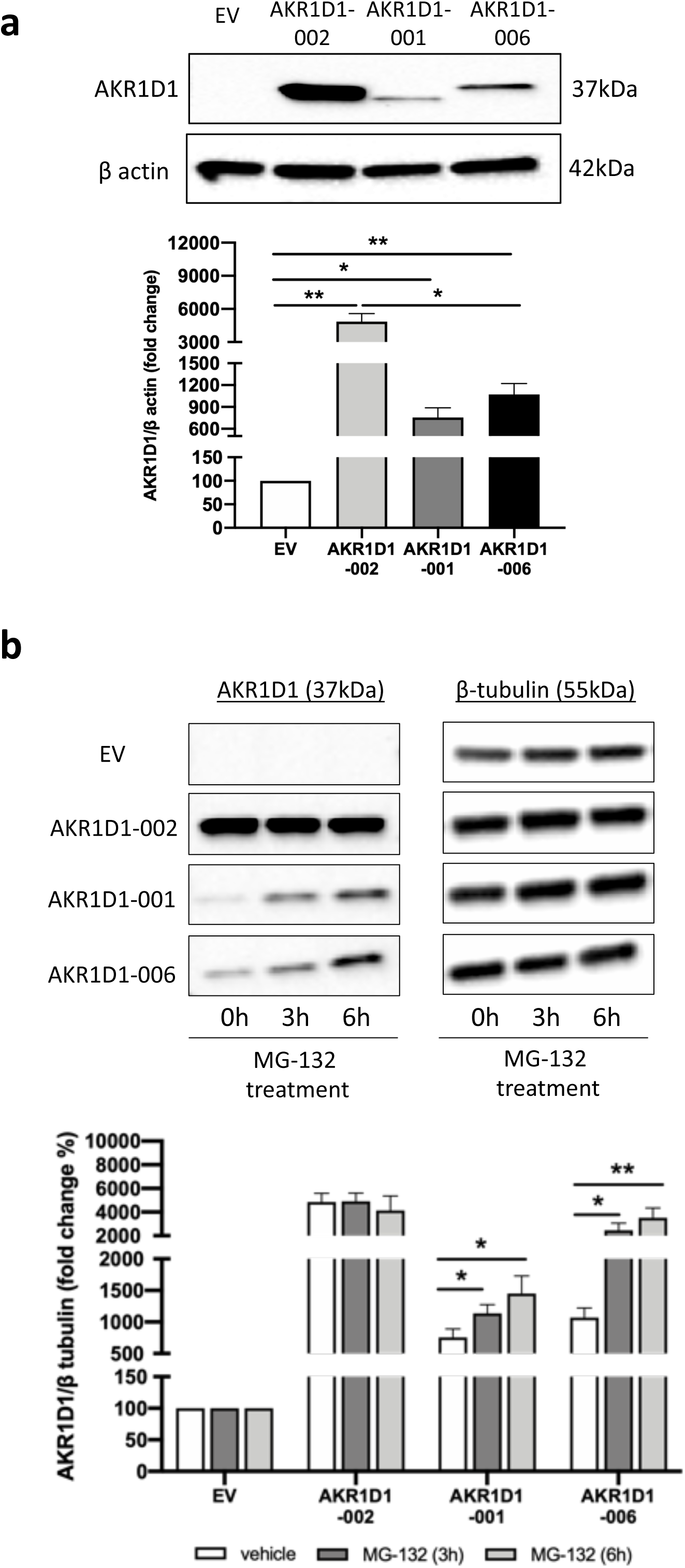
(A) Protein expression levels of AKR1D1-002, AKR1D1-001 and AKR1D1-006, following over-expression in HEK293 cells, as measured by western blotting. (B) Protein expression levels of AKR1D1-002, AKR1D1-001 and AKR1D1-006 following over-expression in HEK293 cells for 48 hours, and subsequent treatment with either DMSO (vehicle, white bars) or 20µM MG-132 (proteasomal inhibitor) for 3 hours (dark grey bars) and 6 hours (light grey bars). Representative western blotting images are shown from 1 biological replicate, however formal quantification was performed in n=4 replicates. Data are presented as mean±se. *p<0.05, **p<0.01, compared to empty vector (EV) transfected controls.

To investigate the hypothesis that truncated *AKR1D1*-SVs undergo proteasomal degradation, HEK293 cells were transfected with either empty pCMV6 (EV), *AKR1D1-002, AKR1D1-001, or AKR1D1-006* containing vectors for 48 hours and treated with the proteasome inhibitor, MG-132 (20μM) for 3 or 6 hours, as previously described (Chui *et al.* 2019). Confirming our hypothesis, MG-132 treatment significantly increased protein levels of both AKR1D1-001 and AKR1D1-006 in a time-dependent manner (Fig. 4B).

### 3.3. *Truncated AKR1D1*-SVs *demonstrate distinct protein structures*

Following *AKR1D1*-SVs over-expression in *in vitro* human cell systems, multiple amino acid sequence alignments were conducted for the three *AKR1D1*-SVs (Clustal Omega) (Goujon *et al.* 2010; Sievers *et al.* 2011). The sequence alignments revealed that the AKR1D1-001 protein misses the amino acids 153-193, whilst the AKR1D1-006 protein lacks amino acids 286-326 and has an additional 5 amino acids at the C-terminus (Val, Ala, Arg, Ser, Ser) (Fig. 5A). Following that, *in silico* protein structure modelling was performed to predict the 3D structures of the truncated variants, using cortisone and NADP+ as substrate and co-factor, respectively (www.PyMol.org). Prediction modelling on AKR1D1-001 revealed that the deleted 153-193 amino acid region could disrupt the interaction between the nicotinamide head of the co-factor and likely hydride transfer to the A-ring of the steroid. In addition, the residues S166 and N170, which bind the carboxamide side chain of the nicotinamide ring, would be absent from the structure (Fig. 5B). In contrast, prediction modelling for AKR1D1-006 demonstrated that the absence of exon 8 plus the addition of five amino acids at the C-terminus lead to the loss of the C-terminal flexible loop (amino acids 286-326), which borders the steroid channel, as well as loss of the last helix in the structure, and is predicted to have decreased affinity for steroid substrates (Fig. 5C).

**Figure 5.**
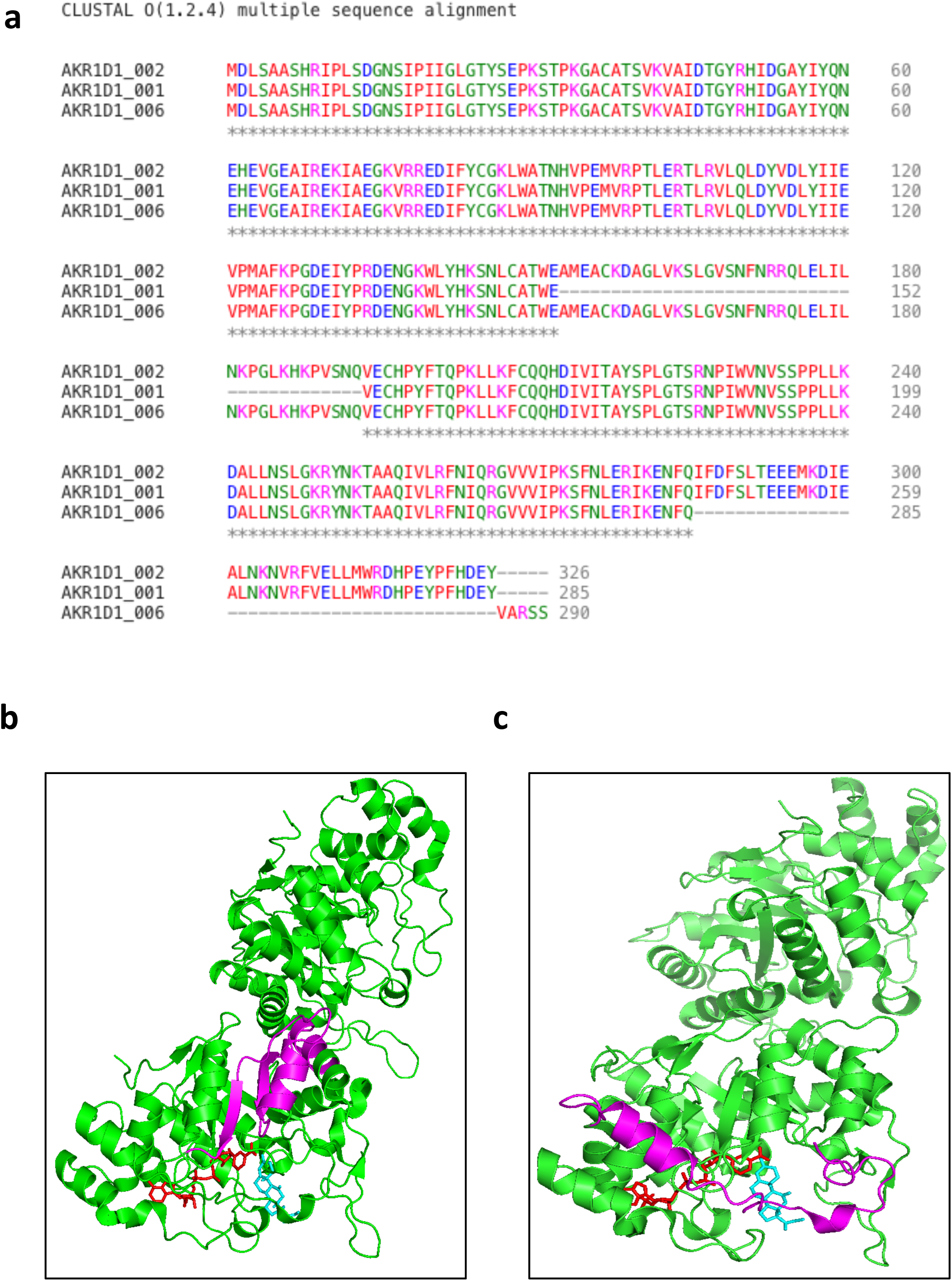
(A) Clustal Omega multiple alignment of the amino acid sequences of the human AKR1D1 splice variants −002, −001 and −006. (B) AKR1D1-001.NADP+.Cortisone Ternary Complex (PDB 3CMF). Magneta shows deletion of residues 153-193; NADP+ (red) and cortisone (blue). There are two monomers per asymmetric unit. (C) AKR1D1-006.NADP+.Cortisone Ternary Complex (PDB 3CMF). Magneta shows deletion of exon 8 (residues 285 – 326); NADP+ (red) and cortisone (blue). There are two monomers per asymmetric unit. Colour coding in figure A represents amino acid physicochemical properties: RED: Small (small + hydrophobic (aromatic)); BLUE: Acidic; MAGENTA: Basic – H; GREEN: Hydroxyl + sulfhydryl + amine + G; ✶: indicates positions which have a single, fully conserved residue. Figures B and C made in PyMol.

### 3.4. *Truncated AKR1D1*-SVs *differentially regulate glucocorticoid and androgen metabolism in vitro*

To investigate the functional activity of truncated *AKR1D1*-SVs, HEK293 cells were transfected with empty pCMV6 (EV), *AKR1D1-002, AKR1D1-001, or AKR1D1-006* containing vectors for 48 hours, and cells were then treated with cortisol, dexamethasone or prednisolone (500nM) for a further 24 hours. There was no change in cell media concentrations for any glucocorticoid in the presence of empty vector (EV). Over-expression of all AKR1D1-SVs significantly reduced cell media dexamethasone concentrations (Fig. 6A). However, only AKR1D1-002 was able to metabolise cortisol and prednisolone (Fig. 6B and C).

**Figure 6.**
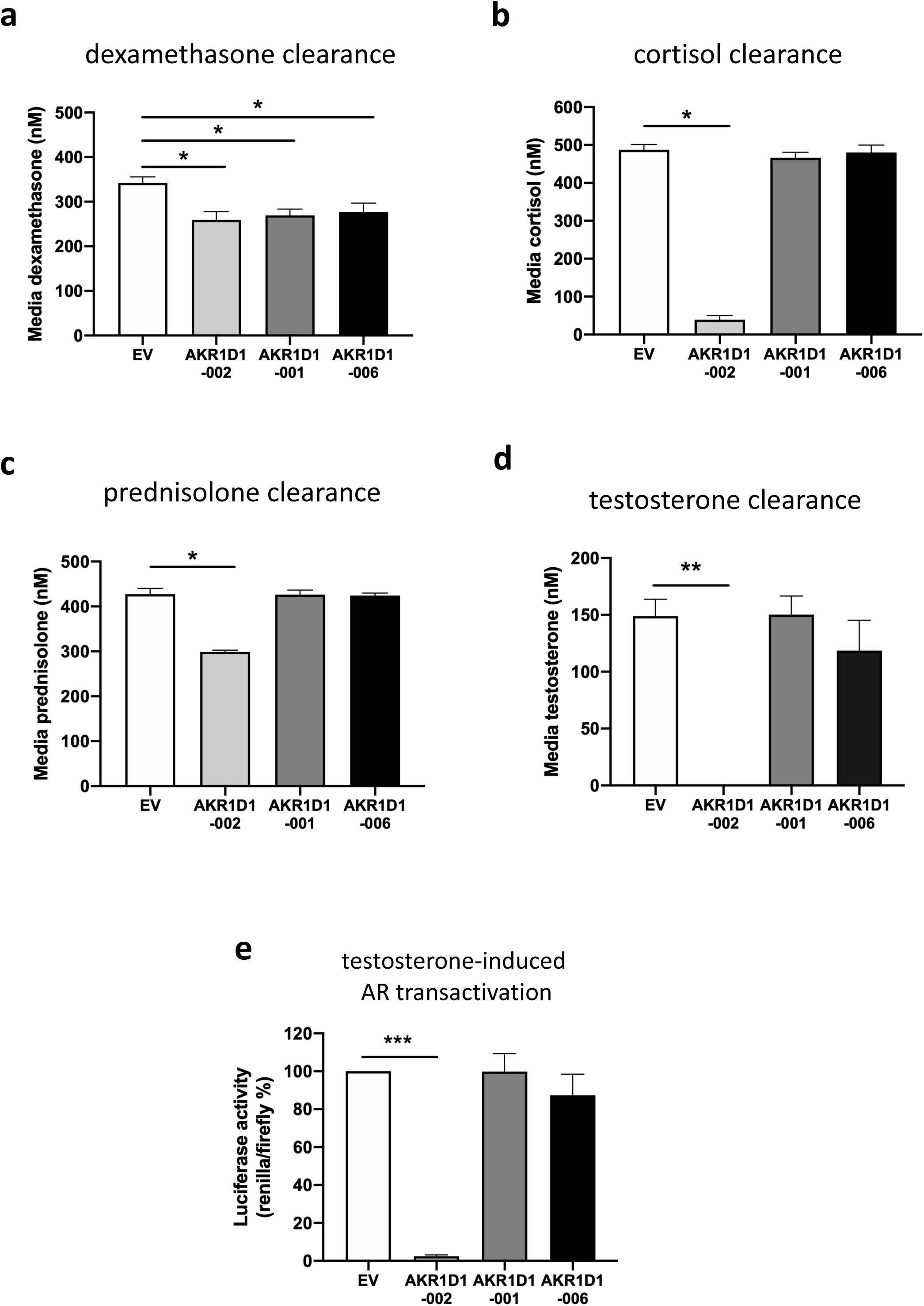
(A) Dexamethasone, (B) cortisol, and (C) prednisolone clearance in AKR1D1-002 (light grey bar), AKR1D1-001 (dark grey bar), AKR1D1-006 (black bar) or empty vector (EV, white bar) transfected HEK293 cells (after 24 hours of steroid treatment, 500nM). (D) Testosterone clearance in AKR1D1-002 (light grey bar), AKR1D1-001 (dark grey bar), AKR1D1-006 (black bar) or empty vector-(EV, white bar) transfected HEK293 cells (after 24 hours of testosterone treatment, 200nM). (E) Testosterone-induced androgen receptor (AR) activation in AKR1D1-002 (light grey bar), AKR1D1-001 (dark grey bar), AKR1D1-006 (black bar) over-expressing and empty vector (EV, white bar) transfected HEK293 cells (white bar) (after 24 hours of testosterone treatment, 200nM). Data are presented as mean±se of n=4 experiments, performed in duplicate, *p<0.05, **p<0.01, compared to empty vector transfected controls.

AKR1D1 is reported to have a crucial role in the regulation of androgen availability (Charbonneau & The 2001). In separate transfection experiments, HEK293 cells were transfected with *AKR1D1*-SV containing vectors or EV (48 hours), and then treated with testosterone (200nM) for 24 hours. AKR1D1-002 over-expression completely cleared testosterone after 24 hours of treatment, however, there was no change in testosterone media concentrations following either AKR1D1-001 or AKR1D1-006 over-expression (Fig. 6D). Consistent with these data, AKR1D1-002 over-expression significantly reduced testosterone-mediated androgen receptor (AR) activation, compared to EV-transfected HEK293 cells. Over-expression of AKR1D1-001 or AKR1D1-006 had no impact on testosterone-mediated AR activation (Fig. 6E).

## 4. Discussion

This study provides the first evidence of the functional role of all *AKR1D1*-SVs in the regulation of glucocorticoid and androgen metabolism. We show that *AKR1D1-001* and *AKR1D1-002* are expressed in human liver biopsies and liver cell lines, whilst *AKR1D1-006* is the predominant variant, and expressed only, in human testes. We demonstrate that, similar to AKR1D1-002, the truncated AKR1D1-001 and AKR1D1-006 proteins metabolise synthetic glucocorticoids (albeit poorly) *in vitro*, but neither of the truncated AKR1D1 proteins metabolise endogenous glucocorticoids and androgens. Finally, we show that truncated *AKR1D1*-SVs undergo rapid intracellular proteasomal degradation.

Examination of the predicted structures of the truncated *AKR1D1*-SVs suggest an effect on function. The exon 5 omitted in AKR1D1-001 does not cause a frameshift in the protein; a previous study from Barski et al. (Barski *et al.* 2013) suggested that, as residues in the middle of the protein sequence are missing, this protein may be structurally compromised, leading to improper folding. Notably the loss of exon 5 will also interfere with the reaction trajectory because S166 and N167 are absent and these stabilise the carboxamide side-chain of the nicotinamide ring (Penning *et al.* 2019). In contrast, AKR1D1-006, which omits exon 8, does cause a frameshift in the protein and misses residues that close over the steroid channel; thus, it is predicted to decrease affinity for steroid substrates (Barski *et al.* 2013). These differences in protein structure are shown in cartoon form in Figure 5B. It is possible that the differences we observed in steroid clearance between the long AKR1D1-002 and the shorter AKR1D1-001/AKR1D1-006 SVs may reflect the structural disruption in the truncated proteins (thus a lower affinity for steroid hormones), or potentially decreased protein stability.

Protein stability has recently been explored in disease-related AKR1D1 mutations (Drury *et al.* 2010). Missense mutations in AKR1D1 have been associated with inherited 5β-reductase and bile acid deficiency, including Leu106Phe, Pro133Arg, Pro198Leu, Gly223Glu, and Arg261Cys1. Following expression in HEK293 cells, these AKR1D1 mutants showed significantly lower protein expression levels than wild-type AKR1D1, despite equal mRNA expression. Analysis of protein degradation rate using a protein synthesis inhibitor, cycloheximide, suggested the mutations impaired protein folding and stability (Drury *et al.* 2010). Importantly, the mutants retained some 5β-reductase activity (*via* detection of 5β-reduced testosterone) despite 100-fold lower expression, indicating the disease phenotypes may not be caused by defects in enzymatic properties, but rather by reduced expression of active AKR1D1 protein.

Similarly, in our study, and despite high mRNA expression, protein levels of both AKR1D1-001 and AKR1D1-006 were significantly lower than those of AKR1D1-002, and proteasomal inhibition treatment partially restored truncated AKR1D1 protein levels. As the ubiquitin-proteasome pathway regulates digestion of misfolded or damaged polypeptides in the cell (Goldberg 2003), it is plausible that omitting exons 5 or 8 in the *AKR1D1* transcripts results in improper post-translational protein folding, therefore degrading the truncated AKR1D1 proteins. AKR1D1 is the first member of the 1D subfamily, along with all known mammalian 5β-reductases, including the rat (AKR1D2), the rabbit (AKR1D3) and the mouse (AKR1D4) homologs (Onishi *et al.* 1991; Faucher *et al.* 2008; Chen *et al.* 2019). Amongst these, two mouse *AKR1D4*-SVs, *AKR1D4L* and *AKR1D4S*, have been recently characterised. Both mouse transcripts were expressed in mouse hepatic and testicular tissues, and enzymatic kinetic assays revealed their ability to metabolise cortisol, progesterone and androstenedione to their 5β-reduced metabolites (Chen *et al.* 2019). Interestingly, they also displayed lower 3α-hydroxysteroid dehydrogenase activity, however, this was limited to C19 steroids, only (Chen *et al.* 2019). In our study, neither of the truncated human *AKR1D1*-SVs metabolised cortisol, testosterone or androstenedione, adding evidence to the presence of distinct functional differences between the human and murine 5β-reductases.

Glucocorticoids are lipophilic molecules that undergo a variety of metabolic conversions to increase their water solubility and enable efficient renal excretion; however, synthetic glucocorticoid clearance has only been examined in a limited number of studies. CYP3A isoforms drive dexamethasone clearance through the formation of 6-hydroxylated metabolites *in vitro* (Gentile *et al.* 1996; Tomlinson *et al.* 1997a, b). In a clinical study, urinary steroid metabolome analysis of samples from healthy male volunteers, following prednisolone administration, identified 20 different prednisolone metabolites, including 11-hydroxylated, 20-reduced and 5α/β-reduced products (Matabosch *et al.* 2015). Supporting these findings, we have recently demonstrated the ability of AKR1D1-002 to clear prednisolone and dexamethasone, with concomitant decreases in hepatic GR activation (Nikolaou *et al.* 2019a, 2020). We now show that, in addition to AKR1D1-002, AKR1D1-001 and AKR1D1-006 metabolise dexamethasone (albeit poorly) *in vitro*. Considering the reduced protein stability of AKR1D1-001 and AKR1D1-006, as well as their inefficient metabolism of cortisol *in vitro*, the role of these truncated *AKR1D1*-SVs in synthetic glucocorticoid clearance is likely to be minimal.

Up to 3% of the UK population are prescribed glucocorticoids therapeutically (predominantly prednisolone and dexamethasone), for the suppression of inflammation in chronic inflammatory diseases, including rheumatoid arthritis, inflammatory bowel disease and asthma (Barnes 1998; van Staa *et al.* 2000; van der Goes *et al.* 2014; Vandewalle *et al.* 2018). In addition, they are used in combination with anti-cancer agents to reduce the genotoxic side effects of chemotherapy treatment, including nausea and vomiting (Collins *et al.* 2007; Buxant *et al.* 2015). Pro-longed use of synthetic glucocorticoids is however associated with adverse metabolic effects including obesity, insulin resistance, and non-alcoholic fatty liver disease (NAFLD) (Woods *et al.* 2015), and decreased AKR1D1 expression has been recently shown in patients with type 2 diabetes and NAFLD (Valanejad *et al.* 2018; Nikolaou *et al.* 2019b). Our study indicates that AKR1D1-002 is the predominant functional variant in human liver; alterations in AKR1D1-002 expression and activity may therefore contribute to the adverse metabolic impact of exogenous glucocorticoids. Nevertheless, the potential additional role of truncated *AKR1D1*-SVs cannot be excluded.

Testosterone is the predominant androgen in males and is synthesised in the Leydig cells in the testis from the precursors dihydro-epiandrosterone (DHEA) and androstenedione (Preslock 1980). Following release into the circulation, the majority of testosterone metabolism (90-95%) to produce inactive androgen metabolites occurs in the liver. AKR1D1-002 is the predominant protein expressed in human liver, and metabolises testosterone *in vitro*, leading to the formation of 5β-reduced metabolites (Kondo *et al.* 1994). Chen at al. (Chen *et al.* 2019) demonstrated low, but detectable levels of *AKR1D4* (the murine homolog of *AKR1D1*) in male mouse testicular tissues, and we now show that *AKR1D1* is expressed in human testes, with *AKR1D1-006* the predominant transcript. Unlike AKR1D1-002, we show that both AKR1D1-001 and AKR1D1-006 fail to metabolise testosterone.

Our study comes with limitations. Over-expression of *AKR1D1* transcripts in HEK293 cells may not accurately reflect physiological expression of the variants *in vivo*. Western blotting of human protein samples would help to determine the actual expression levels of *AKR1D1* splice variants in the liver and the testes. Nonetheless, access to healthy human biopsies is limited, and currently there are no specific AKR1D1 antibodies developed for either AKR1D1-001 or AKR1D1-006 proteins.

In conclusion, we have shown that of the three 5β-reductase (AKR1D1) transcript variants which translate into proteins, *AKR1D1-002* and *AKR1D1-001* are expressed in human liver, and only *AKR1D1-006* is expressed in human testes. AKR1D1-001 and AKR1D1-006 poorly metabolise dexamethasone, but both truncated proteins are intracellularly targeted for proteasome degradation. Additional questions need to be now addressed to dissect the broader role of these variants in translational clinical studies *in vivo.*

## Author contributions

Conceptualisation, N.N.; Methodology, T.M.P., B.K., N.N.; Investigation, N.A., E.G., N.J.D., K.M, S.G., T.M.P., N.N.; Writing - Original draft, N.A., N.N.; Writing - Review & Editing, T.M.P., L.L.G., J.W.T., N.N.; Supervision, J.W.T., N.N.; Funding Acquisition, N.A., J.T.W, N.N..

## Acknowledgments

This work was supported by the Society for Endocrinology (Early Career Grant awarded to N.N., Summer Studentship Grant awarded to N.A.); Medical Research Council, UK (programme grant awarded to J.W.T., ref. MR/P011462/1); NIHR Oxford Biomedical Research Centre (Principal investigator award to J.W.T.), based at Oxford University Hospitals NHS Trust and University of Oxford; P30-ES013508 awarded to T.M.P. by the National Institute of Environmental Health Sciences. The views expressed are those of the author(s) and not necessarily those of the NHS, the NIHR or the Department of Health or the National Institute of Environmental Health Sciences.

